# Ecological niche modeling reveals habitat differentiation and climatic vulnerability in two imperiled, sympatric southern Appalachian carnivorous plants

**DOI:** 10.1101/2025.10.30.685572

**Authors:** Nicholas J. Chang, Lauren Eserman, Amanda Carmichael, Adam B. Smith, Xingwen Loy, Emily E. D. Coffey, James Ojascastro

## Abstract

**PREMISE:** Understanding the habitat requirements of imperiled flora is critical for informing *ex situ* conservation practices, designing effective reintroduction strategies, and understanding how climate change will impact such species, especially in montane regions with high levels of environmental heterogeneity. In southern Appalachia, USA, the Mountain Sweet Pitcher Plant (*Sarracenia rubra* ssp. *jonesii*) and Mountain Purple Pitcher Plant (*Sarracenia purpurea* var. *montana*) inhabit overlapping ranges. These taxa rarely co-occur in the same mountain bogs but frequently hybridize at sites where they do co-occur.

**METHODS:** We assessed patterns of climatic niche differentiation in these imperiled taxa to explore whether they naturally co-occur or may have been brought into secondary contact through human translocations. In addition, we constructed ecological niche models to evaluate the comparative availability of suitable habitat for each taxon under present and future climates.

**RESULTS:** We 1) find evidence that the two taxa inhabit distinct niches, and 2) predict that current populations of *Sarracenia rubra* ssp. *jonesii* will experience climates markedly different from those it currently inhabits in the future, while 3) suitable habitat for *S. purpurea* var. *montana* will remain comparatively stable and may expand in its current range.

**CONCLUSIONS:** Despite high spatial overlap, these two related taxa exhibit divergent climatic niches, resulting in highly different management needs and conservation approaches. We raise concerns about the future of mountain bog plant assemblages, and the rare species they include, under climate change.

## INTRODUCTION

The complex topography of montane regions yields diverse environments that drive evolutionary divergence as organisms adapt to novel environments or become isolated from ancestral populations (Rahbek et al. 2019). In turn, divergence can generate high levels of endemism and rarity (Noroozi et al. 2018, Enquist et al. 2019). Unfortunately, these environments and their unique biodiversity face a variety of threats, including mining (Wickham et al. 2013), urbanization (Terando et al. 2014), and climate change (Potter et al. 2010, Robbins et al. 2024). In particular, elevation-dependent warming may accelerate climate change at higher elevations (Pepin et al. 2015, 2022). Understanding the climatic niche breadth of montane species is key to assessing their vulnerability to climate change and developing effective *in situ* and *ex situ* conservation strategies.

The southern Appalachian Mountains of eastern North America support high biodiversity (Hairston 1949, Estill and Cruzan 2001) and are threatened by climate change (Delcourt and Delcourt 1998, Potter et al. 2010, Milanovich et al. 2010, Ulrey et al. 2016). Among southern Appalachia’s notable habitats are the mountain bogs: isolated, high-elevation wetlands ranging from 0.5–2.0 ha in area and occurring 365–1500 m above sea level (a.s.l.; Richardson and Gibbons 1993). These systems vary widely in their community composition but tend to have high plant diversity, including many rare species (Weakley and Schafale 1994, Edwards et al. 2013), making them priority habitats for conservation (Moffett and Radcliffe 2016, U.S. Fish & Wildlife Service 2021). Among the notable inhabitants of southern Appalachia’s mountain bogs are two highly imperiled carnivorous plants: the Mountain Sweet Pitcher Plant [*Sarracenia rubra* ssp. *jonesii* (Wherry) Wherry, hereafter, SRJ] and the Mountain Purple Pitcher Plant (*Sarracenia purpurea* var. *montana* D.E.Schnell & Determann, hereafter, SPM).

SRJ is currently listed as Endangered under the U.S. Endangered Species Act (U.S. Fish and Wildlife Service 2025), while SPM is currently under review for federal listing status (U.S. Fish and Wildlife Service 2011) and is ranked as Critically Endangered by NatureServe (2025). Despite being congeners, these taxa are distantly related, sharing a common ancestor roughly 4 Mya (Ellison et al. 2012). Both taxa are endemic to mountain bogs in southern Appalachia and the adjacent Piedmont, with each known from approximately 30 extant sites around the Georgia, North Carolina, and South Carolina tripoint (U.S. Fish & Wildlife Service 2021, Weakley and SE Flora Team 2025). These remaining sites are threatened by changes in hydrology, fire suppression, poaching, trampling, wild hogs (*Sus scrofa* L.), and, potentially, hybridization between the two taxa (Moffett and Radcliffe 2016, U.S. Fish & Wildlife Service 2021, Graf et al. 2022).

Despite their largely overlapping ranges (Figure 1) and shared reliance on mountain bog habitats (Weakley and SE Flora Team 2025), the two taxa only occur in the same bogs (i.e., in syntopy) at six known sites, suggesting that some unknown biological or environmental factors may play a role in preventing co-occurrence. Furthermore, in sites where both taxa co-occur, they hybridize to generate the named nothospecies *Sarracenia × charlesmoorei* Mellichamp (2008). Introgression has been detected in both parent taxa (Stephens et al. 2015, Baldwin et al. 2023), suggesting a lack of natural reproductive barriers to support long-term coexistence (Christie et al. 2022). These studies do not address whether samples with evidence for introgression occur at sites where the plants co-occur and do not interpret the time scales across which introgression occurred, which could provide evidence for whether hybridization is natural or the result of human activities.

**Figure 1.**
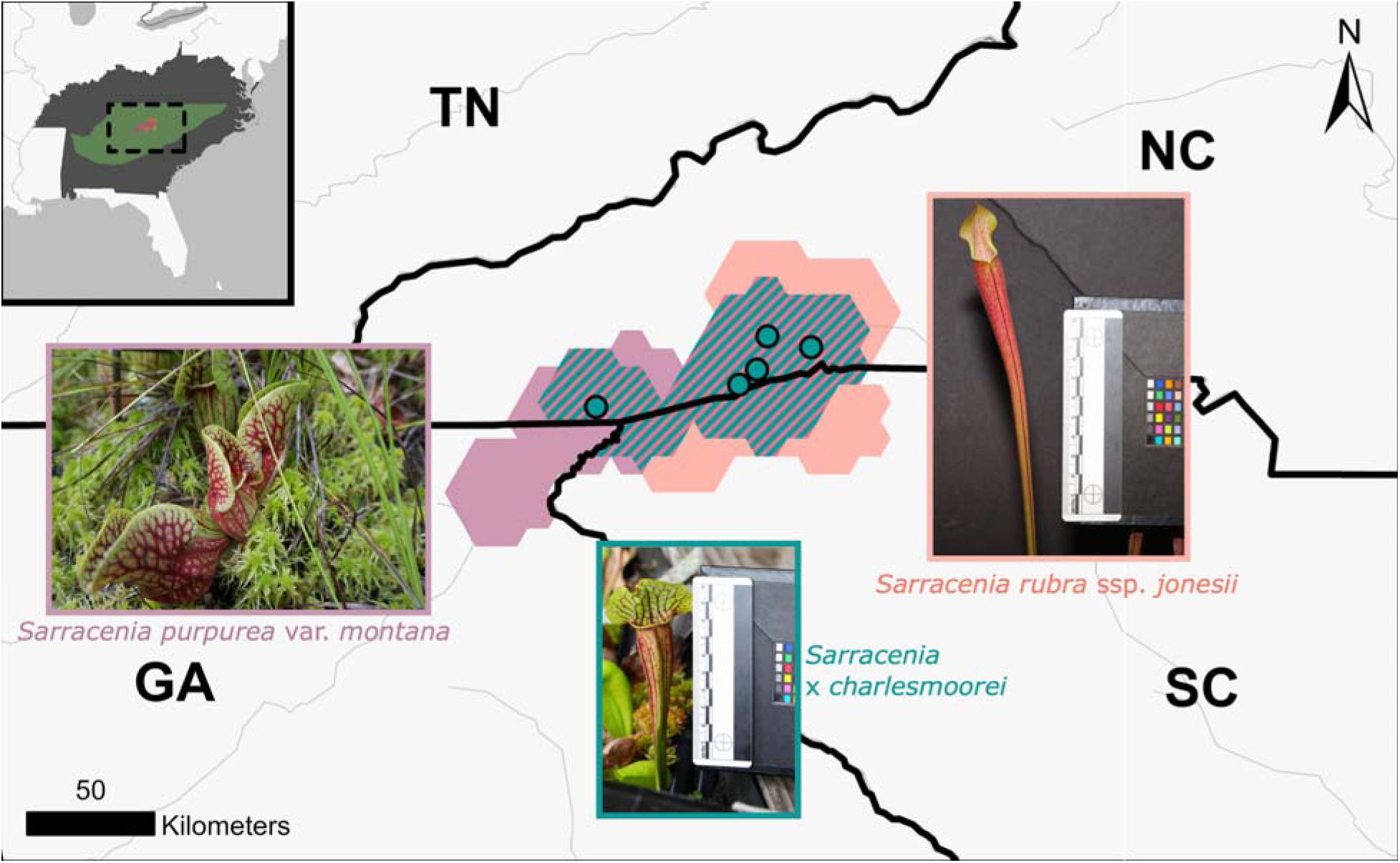
Obscured range extents for *Sarracenia purpurea* var. *montana* (SPM; purple) and *Sarracenia rubra* ssp. *jonesii* (SRJ; red), with overlapping extents shown with teal hatch marks. Known syntopic sites are shown with teal points. Inset map shows the location of the species ranges in the context of the model training area (green) and the area that niche models were projected to (dark gray). Photos courtesy of Nicholas Chang (SPM) and Sathid Pankaew (SRJ and *Sarracenia × charlesmoorei*).

While hybridization can be a natural evolutionary pathway, there are questions about whether some instances of syntopy are the result of historical translocations. These plants have long been the subject of popular interest due to their charisma as carnivorous plants, which makes them both horticulturally interesting and flagship species for mountain bog restoration efforts (Moffett and Radcliffe 2016, Clarke et al. 2018). This attention has resulted in translocation, reintroduction, and population augmentation efforts across several mountain bogs by both private individuals and conservation practitioners (U.S. Fish & Wildlife Service 2021). While herbarium specimens of hybrids date back to at least 1966 (William S. Justice 61: NCU Accession #287803, Catalog # NCU0009244; W.S. Justice 67: NCU Accession #55339, Catalog #NCU00009243), resource managers are concerned that rates of hybridization may be increasing, threatening parental species through genetic swamping and/or a diversion of reproductive investment towards producing hybrid offspring (U.S. Fish & Wildlife Service 2021). As a result, understanding whether current instances of co-occurrence are natural or the result of translocation activities is key to guiding management actions in syntopic sites.

Ecological niche modeling provides a unique framework that allows scientists to inform applied conservation by inferring the climatic variables underpinning both historical and future species distributions. By analyzing the current niches of SRJ and SPM, we can ask whether the two taxa inhabit equivalent or divergent climates. Evidence for niche equivalency between the two taxa might suggest that syntopy was once common, but has declined in recent centuries due to local extirpations and habitat loss. In contrast, evidence for divergence may indicate that these taxa evolved to exploit differing climatic niches, and that current patterns of syntopy are the result of human movement of plant material. This has implications for the management of parent taxa and hybrids in syntopic sites, as well as the planning of future population augmentation efforts. In addition, by using the insights inferred from the present distributions of these plants, we can look to the future and ask how climate change may impact their distributions. The mountain bogs these plants inhabit are small, patchy, and highly dependent on precipitation and temperature regimes, potentially making them extremely vulnerable to climate change; other narrow endemic flora from this region are projected to lose much of their ranges in the future (Schultheis et al. 2010, Erlandson et al. 2021, Hay et al. 2023). Modeling the range dynamics of these taxa under future climates is necessary to select reintroduction sites that will remain suitable habitat in the face of global change (Bellis et al. 2024, 2025).

In this study, we assessed niche divergence and constructed climatic niche models using a dataset of known taxon localities and available climatic raster datasets to evaluate climatic suitability for SPM and SRJ under present and future conditions, with the goal of understanding their present and future range dynamics and informing conservation planning. If both taxa naturally co-occur, we expect no significant difference in their climatic niches and that all six syntopic sites fall within the climatic niche of both taxa. Under a translocation scenario, we would expect divergent climatic niches between taxa, with co-occurrence sites poorly suited to at least one. Intermediate results are plausible, with their interpretation contingent on the nature of interspecific similarities and differences. Finally, we expect that, should these taxa inhabit differing elevational ranges, the higher elevation taxon will undergo a more dramatic range decline under future climate regimes.

## METHODOLOGY

### Data cleaning and processing

We performed all analyses in R version 4.3.1 (R Core Team 2023). We compiled range-wide element occurrence (EO) data for SRJ and SPM from the NatureServe Network (https://www.natureserve.org/). For the purpose of this study, EOs represent all known current and extirpated localities for each taxon that are tracked by state natural heritage programs (NatureServe 2002). We annotated EOs for both taxa based on whether there was prior knowledge that plants had been transplanted. We then overlaid a 30 arc-second resolution raster over these points and manually thinned these occurrences: when multiple occurrences of the same species occurred in the same raster cell, we randomly selected one of the observations to retain and discarded all others; however, when occurrences of both species occurred in the same raster cell, we annotated one observation as “shared” to indicate that both species occurred, and then discarded all other observations from that cell. Given that mountain bogs are small, isolated, and well-documented (Richardson and Gibbons 1993), we assumed that thinning the observations at 30 arc-seconds would result in spatially independent observations without need for further spatial bias correction. To ensure that our presence dataset included the greatest possible number of localities, we supplemented our EO data by requesting redacted coordinate data for *Sarracenia* herbarium specimens from Georgia, North Carolina, and South Carolina (SERNEC Portal Data Portal, 2024). We mapped records of SRJ and SPM onto the same 30 arc-second grid as our EO dataset, and identified two specimens of SPM from North Carolina and one specimen of SRJ from South Carolina with coordinate uncertainties < 1000 meters which did not fall in the same grid cell as an existing EO. We incorporated these herbarium specimens (C. Ritchie Bell 574: NCU Accession #247574; William S. Justice 65: NCU Accession #287806; Robert L. Wilbur #1102: DUKE Accession #126092) into our dataset. We withhold exact locality information to protect these extremely sensitive habitats from trampling (Arnesen 1999) and poaching (Lavorgna and Sajeva 2021); interested parties may request EO data through NatureServe.

We obtained raster datasets of the 19 bioclimatic variables under near-present conditions (1970-2000) at 30 arc-seconds from WorldClim version 2.1 (Fick and Hijmans 2017), and elevation data at ⅓ arc-seconds from the U.S. 3D Elevation Program (U.S. Geological Survey 2024). For projections of future climates, we selected three Coupled Model Intercomparison Project 6 (CMIP6) general circulation models (GCMs) with skillful predictions of eastern North American climates (Almazroui et al. 2021): ACCESS-CM2 (Bi et al. 2013), MPI-ESM1.2 (Gutjahr et al. 2019), and UKESM1-0-LL (Sellar et al. 2019). We note that while UKESM1-0-LL generated skillful predictions of conditions in our study area, globally it has an effective climate sensitivity > 5, suggesting that it may represent higher, “worst-case” future temperature scenarios (Meehl et al. 2020, Hausfather et al. 2022). We obtained downscaled estimates of the 19 bioclimatic variables derived from these GCMs under four Shared Socioeconomic Pathways (SSPs) representing constrained (SSP126 and SSP245), most likely (SSP370), and unabated (SSP585) emissions scenarios at three twenty-year time horizons (2021-2040, 2041-2060, and 2061-2080) and 30 arc-seconds from WorldClim (WorldClim 2022). We averaged across this set of GCMs to obtain a single raster for the mean of each bioclimatic variable at each SSP and time horizon.

### Niche differentiation in environmental space

For analyses in environmental space, we filtered our EOs to remove sites where we had *a priori* evidence for translocation history, and created one subset of EOs that included syntopic sites (SRJ = 29 sites, SPM = 26 sites) and one subset of EOs that excluded these sites (SRJ = 23, SPM = 20). We performed statistical tests on both subsets of data to investigate whether the plants exhibited distinct niches under the assumption that syntopy is a recent phenomenon, and to examine whether our results were robust to the inclusion of syntopic sites. Our first analysis sought to evaluate whether the taxa occur in differing elevational ranges. We extracted elevational data for both sets of EOs, then used a Breusch-Pagan Test to evaluate homoscedasticity and an ANOVA to test for differences in elevation between the two taxa.

Multivariate niche modeling approaches, particularly those trained using sparse occurrence data, are prone to overfitting, prompting us to carefully select a small subset of variables to use in our analyses (Brun et al. 2020). We started with 19 bioclimatic variables (Appendix S1), but excluded elevation, as we sought to identify climatic predictors that have direct physiological implications for these plants, and which may shift along elevational gradients under historical and future climate change scenarios. We extracted values for the 19 bioclimatic variables from all thinned known localities of our focal taxa. This dataset included sites where both co-occur. We then calculated pairwise Spearman’s correlation values for each variable, obtained the absolute value of these correlation values, and used the “hclust” function in the R package *cluster* to perform hierarchical clustering on these variables. In addition, we performed a MULTISPATI, a spatially-explicit ordination method, as implemented in the *R* package *adespatial* version 0.3-28 (Dray et al. 2008, 2025), and inspected the loadings of each of these variables. Using both methods, we selected three bioclimatic variables which were uncorrelated, did not share directions in the ordination, and were biologically interpretable in the context of our study system: annual mean temperature, annual precipitation, and precipitation seasonality (Table 1). Using this set of predictors, we used betadisper() from the *R* package *vegan* version 2.7-2 (Dixon 2003) to test for differences in niche breadth, and performed a PERMANOVA with the *vegan* function adonis(), parameterized using 999 permutations and Euclidean distances, to test for differences in niche occupancy between the focal taxa. We also performed a second MULTISPATI analysis using just the environmental variables retained in the niche differentiation analyses using the R package *adespatial* 0.3-28 (Dray et al. 2025). We regressed the first two axes on elevation to investigate whether observed differences in elevations utilized by these taxa may be driven by climatic differences along mountain slopes.

**Table 1.** Variables retained for use in niche analyses.

| Variable | ANOVA | PERMANOVA | humboldt | MaxEnt |
| --- | --- | --- | --- | --- |
| BIO1 - Annual mean temperature |  | X | X | X |
| BIO7 - Annual temperature range |  | X | X | X |
| BIO12 - Annual precipitation |  | X | X | X |
| BIO15 - Precipitation seasonality |  | X | X | X |
| Elevation | X |  |  |  |

Subsequently, we implemented multivariate tests of niche differentiation in the R package *humboldt* version 1.0.0.0420121 (Brown and Carnaval 2019), where we similarly performed parallel analyses including and excluding syntopic sites. We performed a Niche Overlap Test, which tests for evidence that species occur in different environments across both of their respective ranges. We complemented this analysis with a Niche Divergence Test, which accounts for apparent differences in species niches driven by differing access to analogous habitats (Brown and Carnaval 2019). We performed this analysis with a study area encompassing the Appalachian and Piedmont physiographic provinces within North Carolina, South Carolina, Georgia, Tennessee, and Alabama to sufficiently describe the physiological limits of these species in our analyses by including environmental data from far outside their geographic ranges. We defined background environments for each species as a 100-km buffer around each species’ occurrences, following Simpson and Spalink (2025), and ran each test for 250 iterations.

### Niche modeling in geographic space

For each species, we developed ecological niche models in MaxEnt version 3.4.3 (Phillips et al. 2006) as implemented in the R package *enmSdmX* version 1.2.12 (Smith 2023). We constructed both models using the same set of predictors as were used for PERMANOVA and *humboldt* (Table 1). We trained these models using all known sites of each taxon, thinned to one occurrence per raster cell, including sites where we have *a priori* knowledge of translocation history and where the two taxa co-occur (SRJ = 33 sites, SPM = 33 sites). We trained our models using 10,000 background points randomly sampled from the same study area used in our analyses in *humboldt.* We allowed the algorithm to select an optimal feature class combination, and manually set the regularization multiplier to 10 due to overfitting at lower values.

We evaluated model performance by randomly assigning occurrences to 3 equal-sized groups (k-folds) and testing how well occurrences in two k-folds (the “training” set; 67% of occurrences) predict occurrences in the third k-fold (the “test” set; 33% of occurrences). We repeated this cross-validation procedure twice more to assess mean model performance across all three k-folds. We used three metrics to quantify model performance in our cross-validation: sensitivity, area under the receiver-operating curve (AUC), and Continuous Boyce Index (CBI). Sensitivity is the proportion of predicted presences that were correct (Fielding and Bell 1997).

AUC can be interpreted as the probability that a randomly-drawn presence has a prediction greater than any randomly drawn site within the study region (Mason and Graham 2002). CBI is a score from −1 to 1 that evaluates a model’s ability to predict probability of presence, with a model scoring > 0 performing better than random (Boyce et al. 2002, Hirzel et al. 2006). Since occurrences are assigned to k-folds at random, we bootstrapped the k-fold assignments 100 times and report their averages (Table 3).

To visualize predictions of the distribution of suitable habitat, we projected our model onto rasterized climatic datasets at both present and ensemble future climate datasets. For these predictions, we visualized our rasters onto a study area encompassing the states of Alabama, Georgia, South Carolina, North Carolina, Tennessee, Kentucky, Virginia, and West Virginia to project the availability of suitable habitat outside of our model training area. We extracted suitability values from current known locations and plotted these locations to identify distribution of suitability scores across extant, presumed natural EOs of both taxa (Appendix S2). Based on the distribution of these scores, we demarcated raster cells as suitable (suitability score ≥ 0.6, which encompasses all but one of the extant, presumed natural EOs), marginally suitable (0.6 > suitability score ≥ 0.4), and unsuitable (suitability score < 0.4). For subsequent approximations of Area of Habitat (AOH; Brooks et al. 2019), we extracted the spatial extent of all raster cells with a suitability score > 0.6 using the function “expanse()” in the R package *terra* version 1.7.39 (Hijmans et al. 2023).

## RESULTS

### Niche differentiation in environmental space

SPM inhabits significantly higher elevations than SRJ (Figure 2; excluding syntopic sites: *F_1, 38_* = 32.09, *P* = 1.64×10^-6^; including syntopic sites: *F_1, 53_ =* 19.94, *P* = 4.22×10^-5^). Our elevation data were homoscedastic between both groups (excluding syntopic sites: *X^2^* = 1.324, *df* = 1, *P* = 0.252; including syntopic sites: *X^2^* = 2.673, *df* = 1, *P* = .102). Excluding syntopic and known artificial sites, the elevational range of SRJ spans approximately 310–900 m a.s.l. (mean ± SD = 586 ± 157.07), while the elevational range of SPM spans approximately 540–1100 m a.s.l. (mean ± SD = 842.3 ± 122.63). Using a PERMANOVA and test for homogeneity of multivariate dispersion, we determined that the two taxa occupy distinctive bioclimatic niches (excluding syntopic sites: *pseudo-F* = 37.193, *P* = 0.001, *R^2^* = 0.4757; including syntopic sites: *pseudo-F* = 21.083, *P* = 0.001, *R^2^* = 0.285) but found no difference in niche breadth between the two taxa (excluding syntopic sites: *F* = 0.0716, *P* = 0.7904; including syntopic sites: *F* = 0.3421, *P* = 0.5611). We similarly found that the two taxa exhibit divergent niches using *humboldt* with the Niche Overlap Test (excluding syntopic sites: *Schoener’s D* = 0.16; *P* = 0.004; including syntopic sites: *Schoener’s D* = 0.371, *P* = 0.0319), and this finding held true when the analysis was restricted to analogous accessible environments with the Niche Divergence Test (excluding syntopic sites: *Schoener’s D* = 0.162; *P* = 0.004; including syntopic sites: *Schoener’s D* = 0.366, *P* = 0.02). Background Statistics were non-significant for these tests (Table 2), indicating that we were able to detect niche differences between the two taxa despite both occurring in indistinguishable background environments.

**Figure 2.**
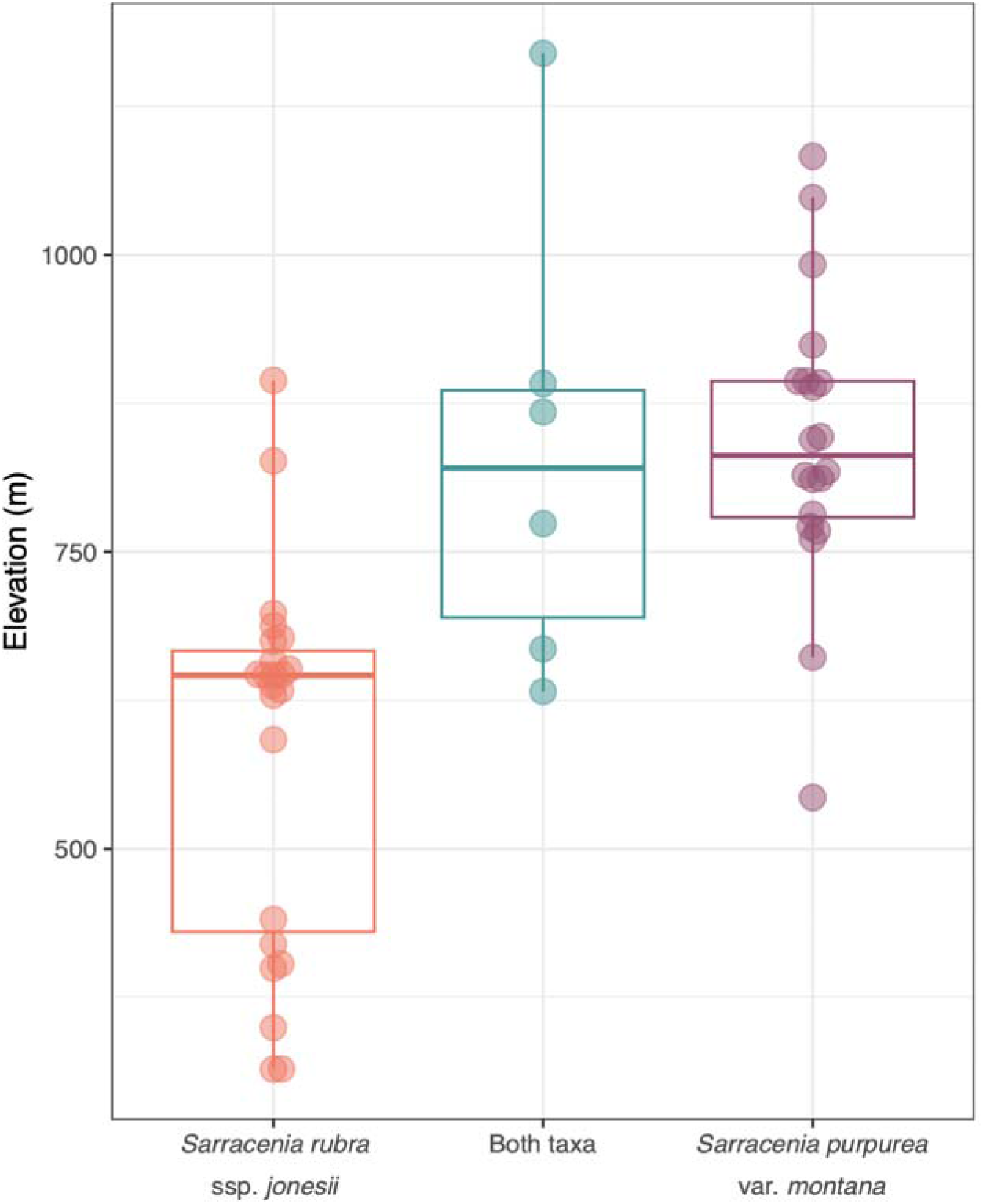
Boxplot showing the elevational distributions of *Sarracenia purpurea* var. *montana* and *Sarracenia rubra* ssp. *jonesii*.

**Table 2.** Outputs of Niche Overlap and Niche Divergence Tests implemented in the R package *humboldt*. *humboldt* incorporates Background Tests that involve shifting the distribution of sites for Taxon 1 and comparing the resulting environmental niche to Taxon 2, and vice versa (Brown and Carnaval 2019). Taxon 1: *Sarracenia rubra* ssp. *jonesii*; Taxon 2: *Sarracenia purpurea* var. *montana*.

|  | Equivalency Test |  | Background Test |  |
| --- | --- | --- | --- | --- |
|  | Schoener's D | P-value | 2 → 1 | 1 → 2 |
| Niche Overlap Test (excluding syntopic sites) | 0.16 | 0.004 | 0.331 | 0.155 |
| Niche Overlap Test (including syntopic sites) | 0.371 | 0.0319 | 0.068 | 0.076 |
| Niche Divergence Test (excluding syntopic sites) | 0.162 | 0.004 | 0.402 | 0.195 |
| Niche Divergence Test (including syntopic sites) | 0.366 | 0.02 | 0.059 | 0.084 |

MULTISPATI analysis suggests that differences in climatic niche between the two taxa are driven by differences in annual precipitation and annual mean temperature (Figure 3). The first axis explained 61.03% of the variation, while the second axis explained 30.74% of the variation, and both MULTISPATI axes were strongly correlated with elevation (Axis 1: β ± SD = −0.0059 ± 0.00057, *Adjusted R^2^* = 0.694, *F_1,47_* = 109.9, *P* = 6.82×10^-14^; Axis 2: β ± SD = 0.003 ± 0.00035, *Adjusted R^2^* = 0.619, *F_1,47_* = 79.02, *P* = 1.24×10^-11^). Three sites containing both taxa are positioned in a cluster of points containing both SRJ and SPM. An additional two sites cluster with SRJ sites, and a final site is closest in environmental space to SPM sites, but is placed outside of the visualized point cloud for all other sites. This final, most divergent site has the highest elevation of any known locality for these taxa (elevation > 1200 m a.s.l.).

**Figure 3.**
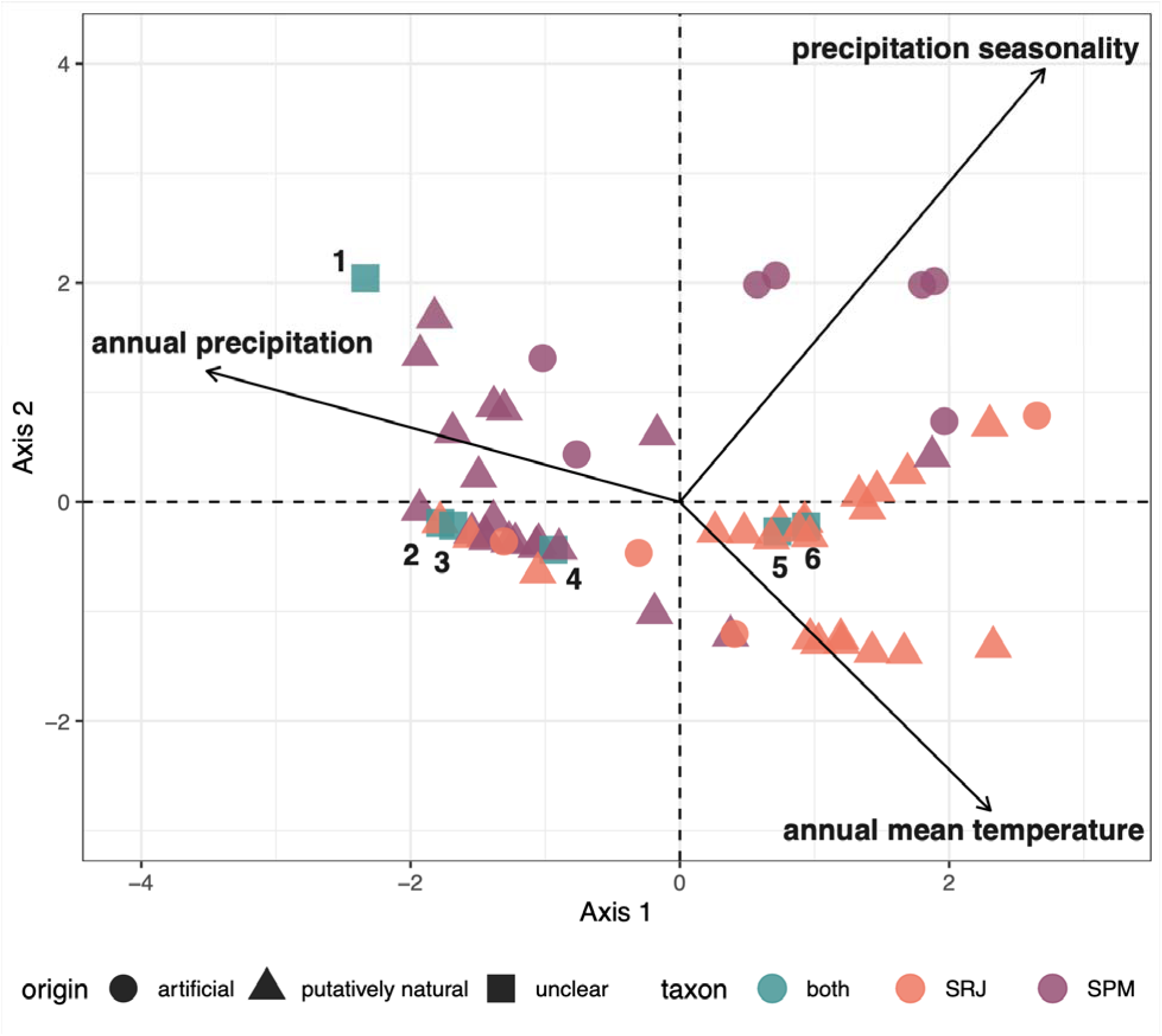
Multivariate spatial analysis based on Moran’s I (MULTISPATI) of bioclimatic environments inhabited by *Sarracenia rubra* ssp. *jonesii* (red), *Sarracenia purpurea* var. *montana* (purple), or both taxa (blue, numbered).

### Niche modeling in geographic space

The MaxEnt models performed well for both species (sensitivity ≥ 0.366, AUC ≥ 0.955, and CBI ≥ 0.827; Table 3). Across the region, suitable habitat for SRJ was concentrated in its current range, as well as small outlying areas in the Allegheny Mountains and in south-central Virginia far from its known distribution. In contrast, suitable habitat for SPM was located exclusively within its current range (Figure 4; Appendix S3). Under most scenarios, both taxa appear to exhibit declines in areas modeled as suitable under the 2021-2040 time horizon, followed by moderate gains during the 2041-2060 time horizon, and then losses again at the 2061-2080 time horizon. Both focal taxa experienced qualitatively similar changes in the proportion of suitable habitat when using a relaxed suitability threshold (Appendix S4).

**Table 3.**
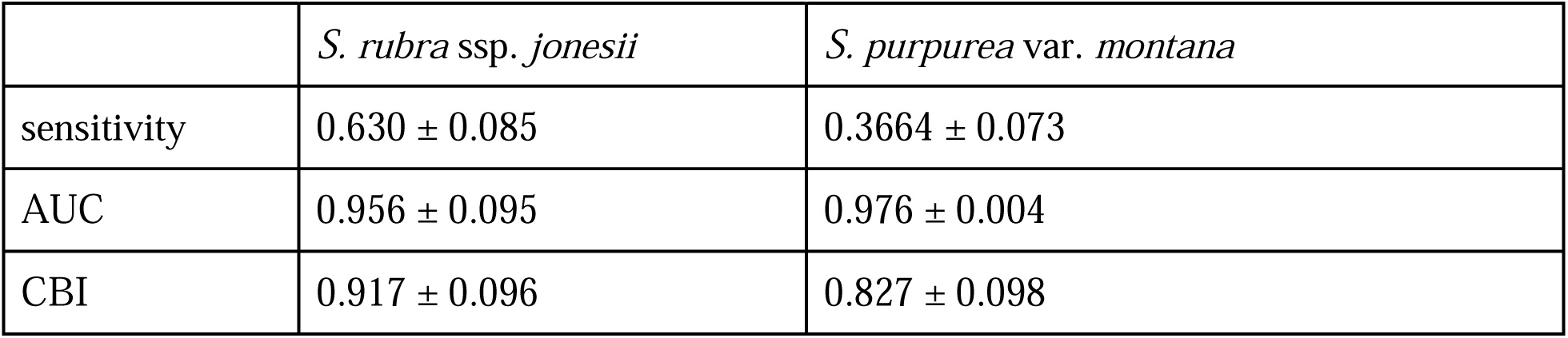
Mean performance of 100 bootstrapped MaxEnt models generated for each taxon using 3 k-folds, ± standard deviation.

|  | *S. rubra* ssp. *jonesii* | *S. purpurea* var. *montana* |
| --- | --- | --- |
| sensitivity | 0.630 ± 0.085 | 0.3664 ± 0.073 |
| AUC | 0.956 ± 0.095 | 0.976 ± 0.004 |
| CBI | 0.917 ± 0.096 | 0.827 ± 0.098 |

**Figure 4.**
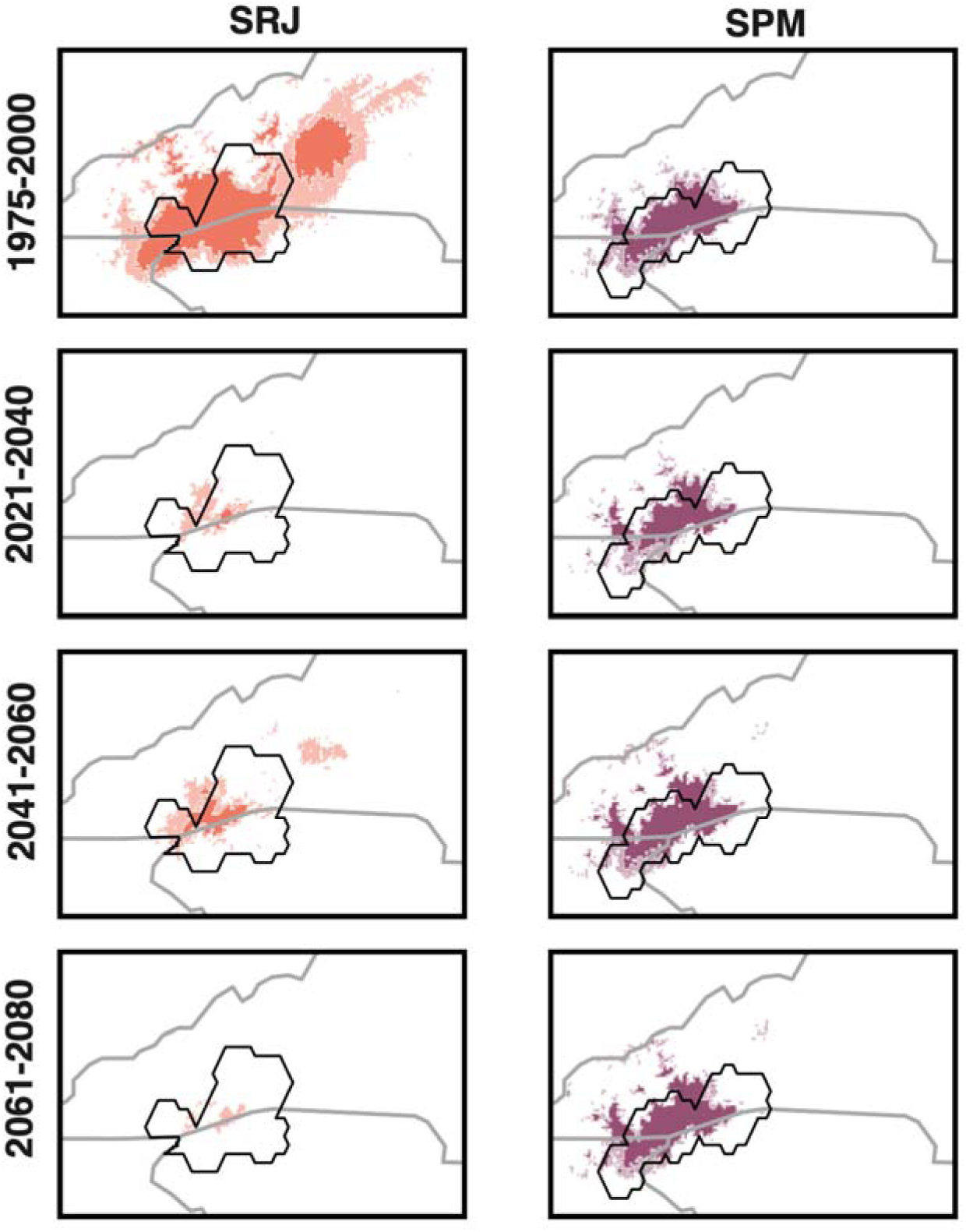
Suitability maps for *Sarracenia rubra* ssp. *jonesii* (SRJ) and *Sarracenia purpurea* var. *montana* (SPM) at near-present and three future time horizons (2021-2040, 2041-2060, 2061-2080) under the most likely emissions reduction scenario (SSP 370). Darker colors indicate areas with suitability scores > 0.6, while lighter shaded areas indicate areas with suitability scores < 0.6 and > 0.4. Black outlines show the actual range extents of each taxon. Full-extent maps, including spatially disjunct areas of projected future suitability north of the current range, are provided in Appendix S3; these broader-scale projections follow the same pattern of declining suitability within the core range shown here.

Models predicted a drastic decline in habitat area within the area occupied by the current climatic niche of SRJ across all SSPs and future time horizons. Under SSP 370, our models estimate that Area of Habitat for SRJ will decline by 95% by 2080 (Figure 5). Of 16 presumed natural, extant sites with near-present suitability scores > 0.6, none will have scores > 0.6 and only 3 of these sites will have scores > 0.4 by 2080 (Figure 6). Under future climates, our models predict limited habitat for SRJ remaining in its current range under the 2021-2040 and 2041-2060 time horizons, but such habitat will vanish by 2080. By 2080, the outlying area in the Allegheny Mountains identified above is the only region in our study area predicted to contain habitat matching the near-present climatic niche of this taxon.

**Figure 5.**
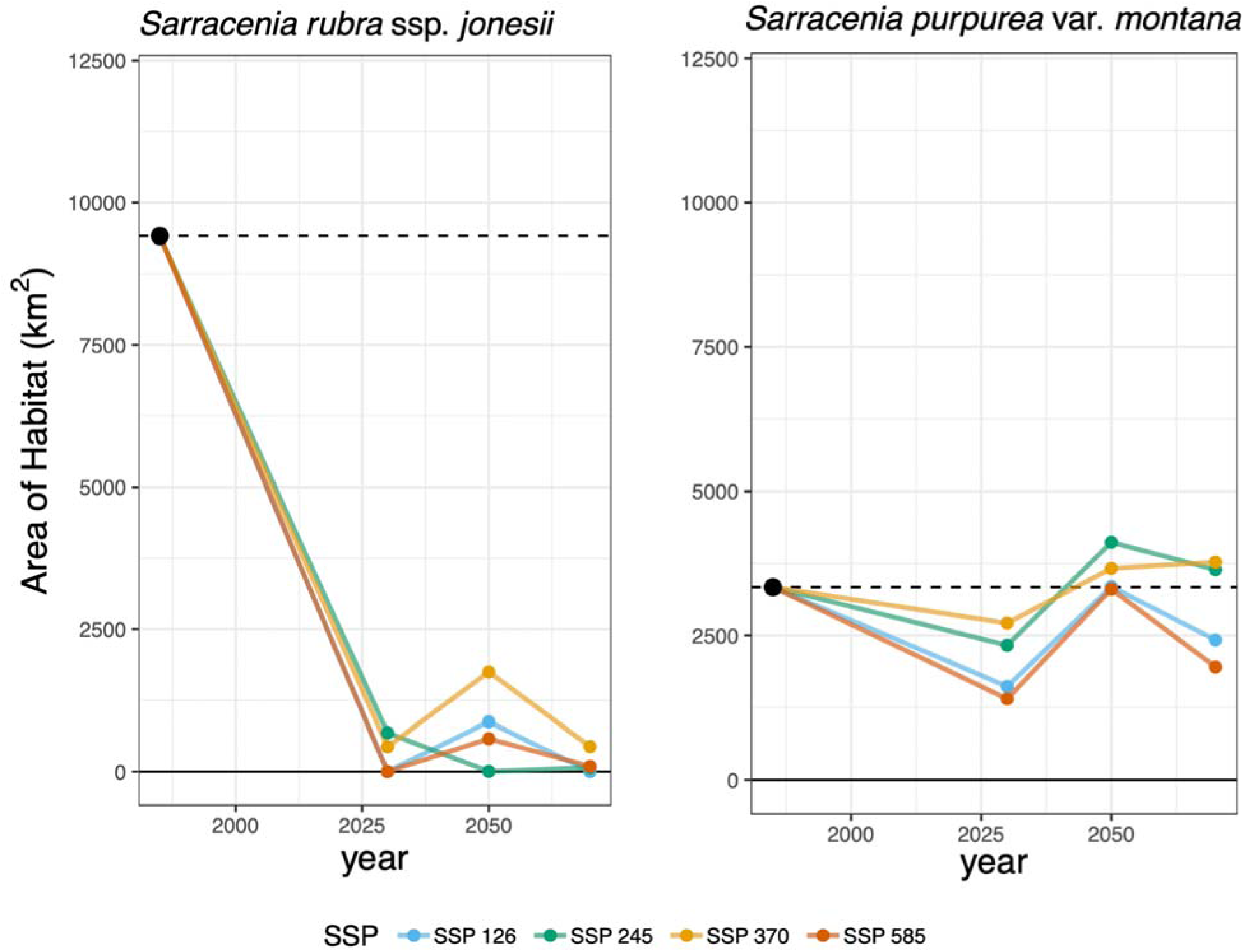
Line graph showing the projected Area of Habitat (AOH; area of raster cells with suitability scores > 0.6) for both taxa at all time horizons and SSPs. Points are mapped to the midpoint of the time horizon modeled (i.e., the point for 2021-2040 is placed at 2030).

**Figure 6.**
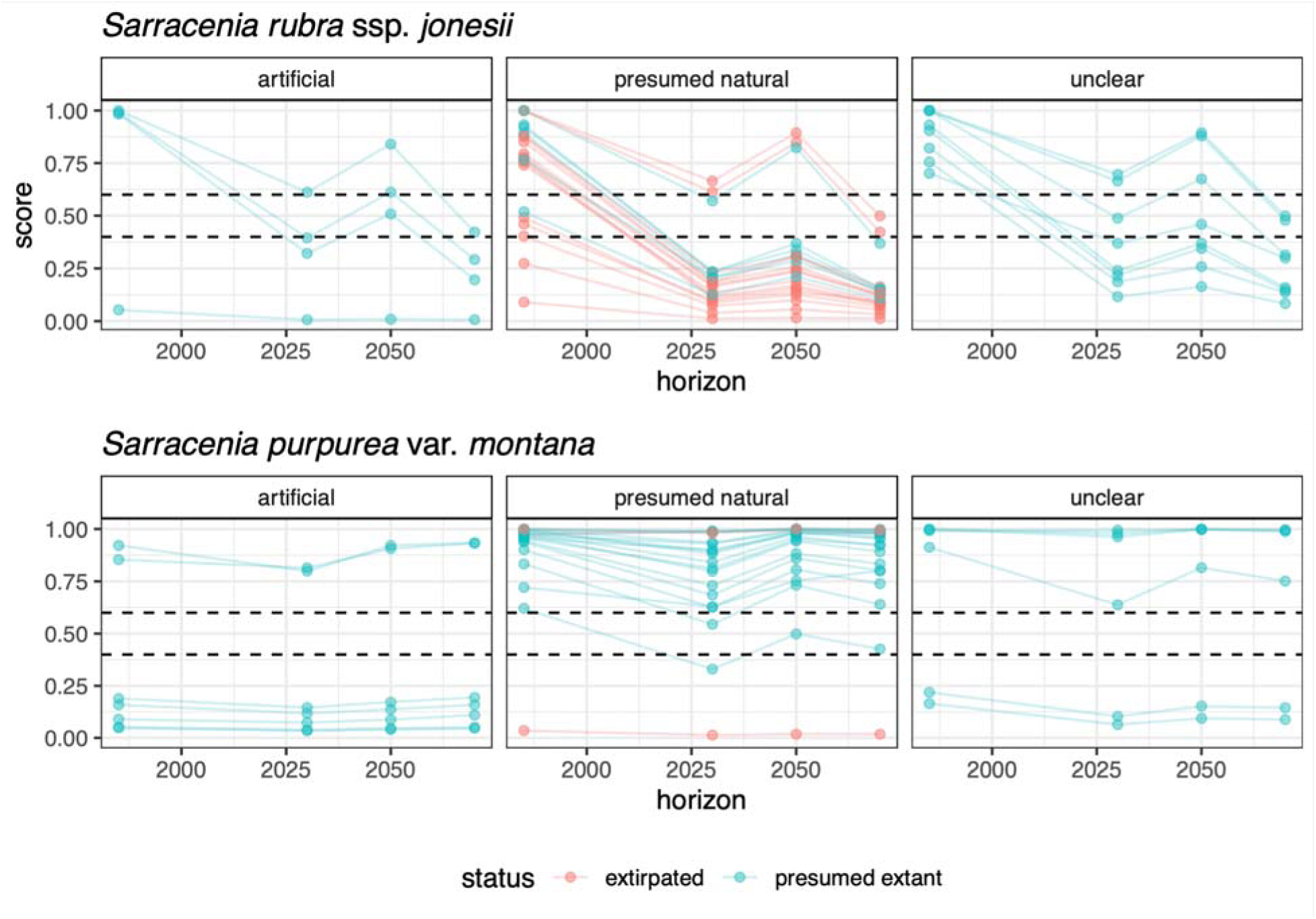
Suitability scores generated by MaxEnt for each site for *Sarracenia rubra* ssp. *jonesii* and *Sarracenia purpurea* var. *montana* under near-present and projected future climates under SSP 370. Dashed lines indicate suitability thresholds at 0.6 and 0.4.

In contrast, models predict that suitable habitat for SPM will remain relatively stable, and may even marginally increase under certain scenarios. We estimate that the Area of Habitat for SPM will increase by 13% by 2080 under SSP 370 (Figure 5). Of 22 presumed natural, extant sites that have near-present suitability scores > 0.6, all 22 sites retained scores > 0.6 by 2080 (Figure 6). Under present and future conditions, we predict that only very minimal areas outside of its immediate geographic range match its present climatic niche, and do not predict a pronounced latitudinal shift in habitat matching the niche of this taxon.

## DISCUSSION

### Contemporary niche differentiation

We found that despite having overlapping geographic ranges, two narrowly-endemic pitcher plants inhabit largely distinct, elevation-correlated climatic niches, indicating that range-wide syntopy is uncommon. Specifically, only three of sixty total sites (5%) occupied by either species fall within climatic space predicted to support both taxa. This interpretation is supported by the low frequency of co-occurrence across all known sites, as well as reports from conservation horticulture staff at the Atlanta Botanical Garden, where SRJ thrives in warm greenhouse conditions alongside Coastal Plain *Sarracenia,* while SPM performs better in cooler, climate-controlled conditions (John Evans, pers. comm.).

Of the six known syntopic sites, three occur within climatic space independently predicted to support both taxa, consistent with limited natural overlap at ecotonal or elevational boundaries. The remaining three sites cluster with one of the two species in our ordination.

Persistence of both taxa at these locations suggest that local microclimate, non-climatic factors, or model uncertainty due to sparse occurrences may allow coexistence beyond modeled climatic envelopes. Thus, while modeled niche overlap is minimal, these results highlight both the rarity of syntopy at broad scales and the limitations of coarse-resolution climate models for interpreting coexistence at individual sites.

Even if co-occurrence occurs naturally, they likely represent rare, peripheral events in each taxon’s evolutionary trajectory that should not necessarily be promoted in conservation practice due to the lack of reproductive barriers in this system and the still-unknown implications of hybridization for the conservation of these two taxa. While hybridization can be an important adaptive pathway that can confer climate-adaptive alleles to imperiled taxa (Brauer et al. 2023), unchecked hybridization between taxa without reproductive barriers can result in extinction by genetic swamping (Todesco et al. 2016). Genetic analyses are needed to determine whether human-mediated translocations underlie current patterns of syntopy and to assess the extent, timing, and origin of hybridization in this system.

As conservation practitioners recognize that the climatic suitability of recipient outplanting sites is an important predictor of translocation success (Bellis et al. 2020, 2025), our findings emphasize the need to understand a species’ niche on a fine spatial scale, particularly in environmentally heterogeneous montane landscapes, to appropriately inform outplanting strategies. The niche of one taxon may not necessarily be useful as a proxy for the niche of another related taxon. Elevational ranges may be helpful for guiding outplanting locations (SRJ: 310–900 m; SPM: 540–1090 m), though for SRJ in particular, habitat suitability under future climate regimes should be considered. As finer-resolution geospatial datasets become available, niche modeling tools will become increasingly useful in assisting conservation practitioners in appropriately designing management and reintroduction strategies.

### Climate change vulnerability

We predicted a major decline in habitat matching the current climatic niche of SRJ under all future climate scenarios, including in the very near term during the 2021-2040 time horizon. In contrast, we predicted that SPM may experience some decline in habitat, but will maintain a relatively stable range compared to SRJ. This finding contrasted with our expectations, given that SPM is associated with higher-elevation habitats than SRJ.

While we predict that suitable habitat for SRJ may occur in more northern areas of Appalachia (i.e. the Allegheny Mountains) under future climates, this region is approximately 450 km north of its current range. *Sarracenia* seeds are water-dispersed, and poleward migration under this mechanism would require SRJ’s seeds to travel hundreds of kilometers downstream through the Tennessee River and then upstream into the Ohio River, which is highly improbable (Ellison and Parker 2002, Zellmer et al. 2012). Some natural mechanisms of rare, long-distance dispersal, such as by migratory animals, likely exist, though the exact mechanisms and frequency of such events are still unknown (Ellison and Parker 2002). However, the Allegheny Mountains also support existing populations of *Sarracenia purpurea* L. ssp. *purpurea,* with which SRJ can hybridize. While assisted migration may warrant serious consideration to avoid the extinction of some southern Appalachian plant species such as *Shortia galacifolia* Torr. & A. Gray (Diapensiaceae)*, Phacelia fimbriata* Michx. (Hydrophyllaceae), and *Gymnocarpium appalachianum* Pryer & Haufler (Cystopteridaceae; Erlandson et al. 2021, Hay et al. 2023), practitioners must carefully weigh the ecological risks and benefits of such long-distance translocations while prioritizing *ex situ* germplasm conservation as a proactive measure against future habitat loss (Browne et al. 2024), following *ex situ* conservation best practices (Guerrant et al. 2010). In contrast, our models predict relatively stable suitable habitat for SPM centered around the Georgia-North Carolina-South Carolina tripoint under all future scenarios. This narrow range suggests adaptation to specific and regionally-unique climatic regimes that may be resilient to past and projected climate shifts. Conservation should prioritize increasing the resiliency and redundancy of populations within its current range extent and adjacent suitable areas.

Similar to other research on the climate futures of southern Appalachian flora, we predicted declines in suitable habitat area in the immediate near term (2021-2040) and towards the end of the century (2061-2080), but moderate gains in suitable habitat in the middle of the century (2041-2060; Potter et al. 2010). These predictions held true among three of the four SSPs forecasted for each taxon. Counter to our expectations, our models did not predict that the severity of a given SSP corresponds with the severity of changes in habitat suitability for a given taxon under that SSP. For example, in SRJ, we predicted that by the middle of the century, the taxon may lose all suitable habitat under SSP 245 (a lower emission climate scenario), while the more moderate scenario, SSP 370, predicted the greatest amount of suitable habitat for that taxon at that time horizon. This mismatch may be driven by regional heterogeneity in climate impacts, and the global severity of an emissions reduction pathway may not correspond to its realized impacts on specific, narrow-ranged taxa. Future assessments of climate futures for rare species may consider increasing the number of GCMs examined and projecting trained niche models onto each GCM individually rather than ensembling GCMs beforehand, to assess variability in predicted future range sizes across each GCM.

While this study provides valuable insights on potential historic and future range dynamics of SRJ and SPM, it has some limitations. Our models use current distributions of both taxa, which have been altered by recent centuries of land-use change that curtailed the potential full extent of inhabited climates. Additionally, correlative models do not necessarily capture all of the biological realities of climate change, and there is a need for studies examining specific drivers of climate vulnerability. Controlled experiments manipulating water availability and temperature, alongside detailed niche studies that quantify factors such as pollinator interactions, stress tolerance, dispersal ability, and soil seed bank longevity, may be needed to clarify species’ responses to changing conditions and their potential for long-term coexistence in syntopic sites. Furthermore, our analyses do not account for the acute impacts of climate-driven extreme weather events (Bateman et al. 2012). During Hurricane Helene in 2024, at least one population of SRJ was physically dislodged from a cataract bog and would have been lost without intervention from the South Carolina Department of Natural Resources and Bartlett Tree Laboratories and Arboretum (Adam Black, pers. comm.). The narrow ranges and specific habitats of these taxa may render them more vulnerable to extreme weather events and may therefore be more threatened by climate change than our models indicate.

Future research is needed to determine the extent to which SRJ faces an imminent range contraction and to ensure that such information informs upcoming conservation assessments and recovery planning. In contrast, conservation efforts for SPM should focus on maintaining and restoring existing habitat within its current range until more detailed recovery plans can be developed. Both taxa would benefit from *ex situ* collections based on maternal line best practices for promoting genetic representation (Guerrant et al. 2010).

This study also highlights the need for improved, region-wide monitoring and management of southern Appalachian bogs. Though scarce, records of bog community composition show considerable turnover over the past five decades, including species extirpations and immigrations (J. Ojascastro, unpubl. data). Comprehensive, geo-referenced records of local floristics, management actions, and species translocations are essential for anticipating future crises, evaluating the impacts of interventions, and enabling the efficient and effective conservation of Appalachian biodiversity.

## Conclusion

Under present conditions, SRJ and SPM inhabit distinct climatic niches, potentially driven by differences in elevation. Until genomic evidence identifies whether hybridization and syntopy are natural in this system, we recommend that resource managers avoid reintroducing one taxon into a site already occupied by another. Additionally, our models suggest SRJ may experience a drastic loss of suitable habitat under future climate regimes, while SPM is expected to remain comparatively secure. *Ex situ* collections development is needed to secure range-wide germplasm from both taxa. Beyond *Sarracenia,* our findings provide a troubling preliminary indication for further range declines in certain Appalachian bog flora. Future research, using geospatial, observational, and experimental approaches, is urgently needed to understand how mountain bog species and assemblages will respond to the anticipated and already-realized effects of climate change. Such work will be key to assisting conservation practitioners and policymakers in charting a new path forward towards conserving and restoring these unique, fragile habitats.

## ACKNOWLEDGEMENTS

The authors thank the editors and reviewers for their insightful comments that greatly improved the quality of this work. This work was funded by the National Science Foundation (Award #2216650), the Atlanta Botanical Garden, and the University of Minnesota. The authors thank the South Carolina and Georgia Departments of Natural Resources and the North Carolina Plant Conservation Program for sharing the element occurrence data analyzed in this work. The authors thank Carrie Radcliffe and Quix Malpass for sharing helpful, site-specific insights, and Milo Vasquez for his administrative support. The authors humbly acknowledge the Cherokee people, who have stewarded and produced knowledge on the lands inhabited by *Sarracenia rubra* ssp. *jonesii* and *S. purpurea* var. *montana* since time immemorial. Data analysis and the writing of this manuscript took place in Atlanta, GA, USA on the traditional homelands of the Mvskoke (Creek) people and in Saint Paul, MN, USA on the traditional homelands of the Dakota people.

## AUTHOR CONTRIBUTIONS

**Nicholas Chang:** conceptualization, methodology, formal analysis, data curation, writing – original draft, writing – review & editing. **Lauren Eserman:** funding acquisition, conceptualization, project administration, resources, writing – review & editing. **Amanda Carmichael:** conceptualization, project administration, resources, writing – review & editing. **Adam Smith:** methodology, software, writing – review & editing. **Xingwen Loy:** writing – review & editing. **Emily Coffey:** funding acquisition, data curation, resources, writing – review & editing. **James Ojascastro:** project administration, conceptualization, resources, methodology, software, formal analysis, writing – review & editing.

## DATA AVAILABILITY STATEMENT

These plants are highly vulnerable to poaching for the horticultural trade, and their habitats are highly sensitive to trampling. As a result, we decline to share the coordinates used to extract bioclimatic data, as well as the extracted bioclimatic data, which could be used to reverse-engineer species locations. Interested parties are encouraged to contact NatureServe for data access. Additional supporting information may be found online in the Supporting Information section at the end of the article.

