## Supplemental Information for "Ecological niche modeling reveals habitat differentiation and climatic vulnerability in two imperiled, sympatric southern Appalachian carnivorous plants"

### Online Supporting Information

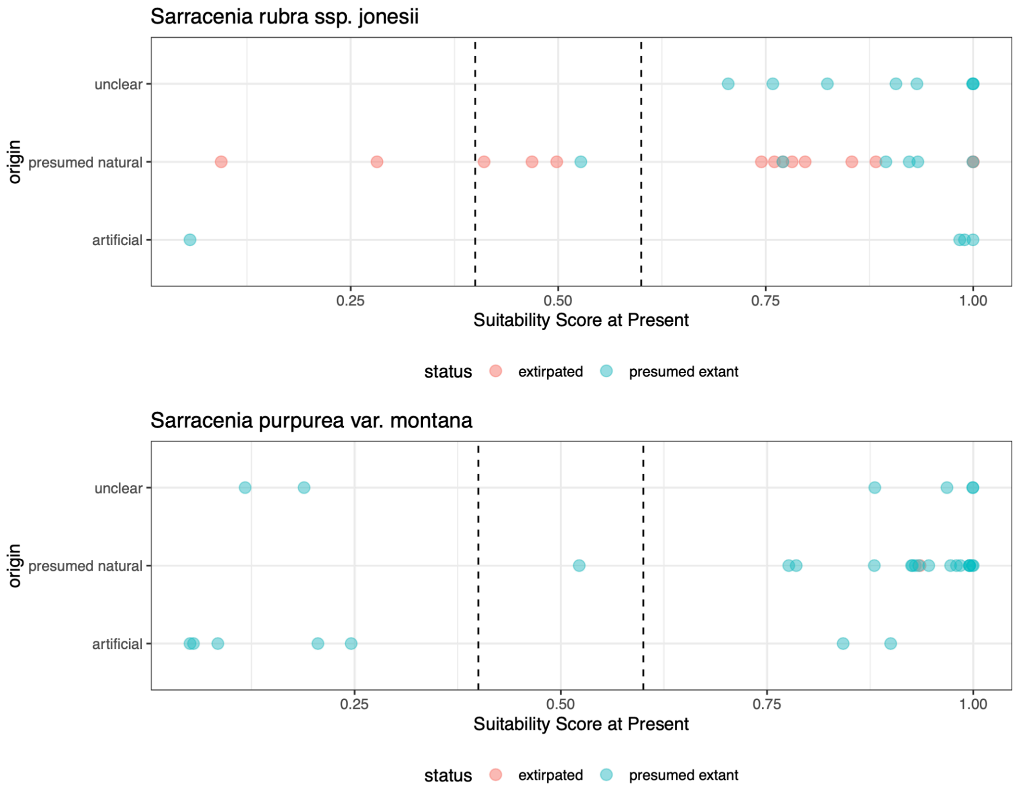


Appendix S1. Historical suitability scores for sites containing either *S. rubra* ssp. *jonesii* or *S. purpurea* var. *montana.*

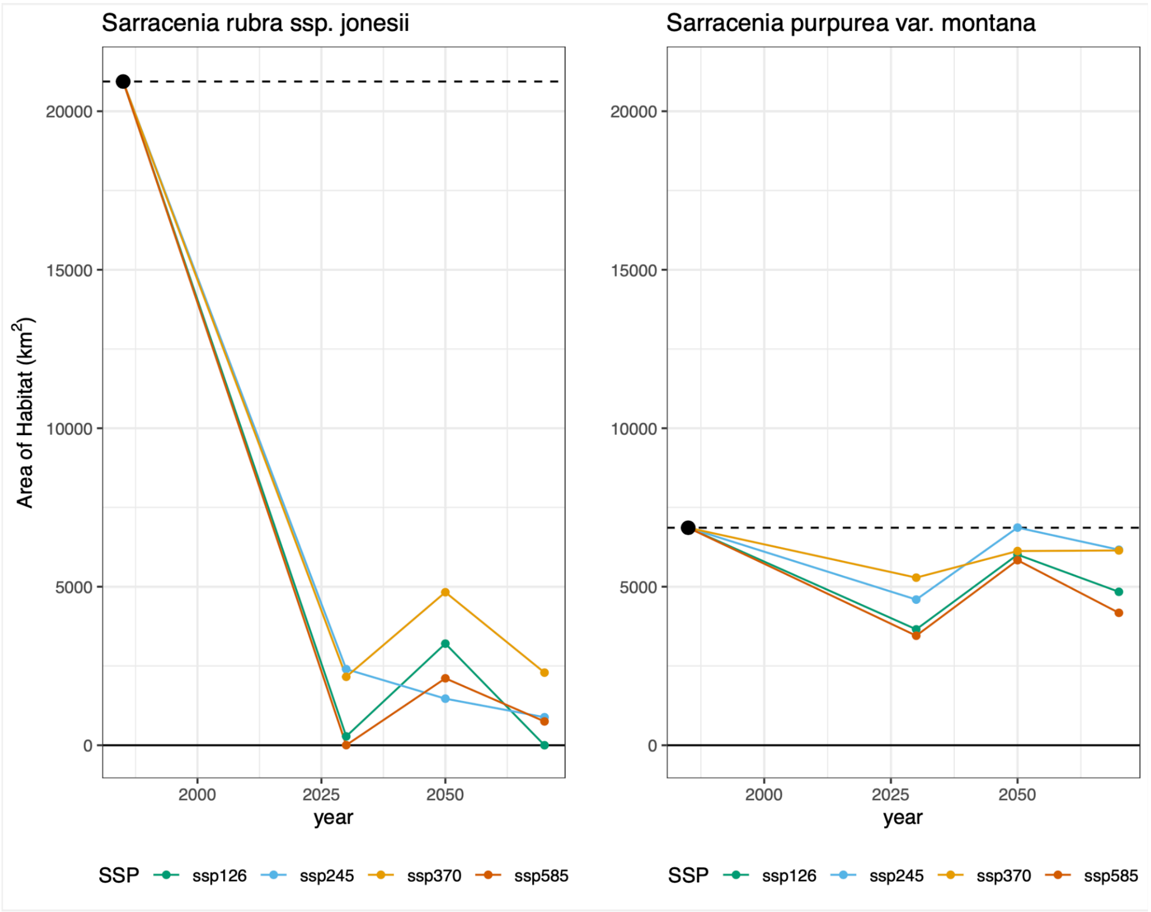


Appendix S2. Projected estimates for Area of Habitat (AOH) of each taxon across all time horizons and SSPs, for areas with a suitability score > 0.4.
